# mol2sphere: Spherical Decomposition of Multi-Domain Molecules for Visualization and Coarse Grained Spatial Modeling

**DOI:** 10.1101/257857

**Authors:** Joseph Masison, Paul J. Michalski, Leslie M. Loew, Adam D. Schuyler

## Abstract

**Summary:** Proteins, especially those involved in signaling pathways are composed of functional modules consisting of long strings of amino acids. These functional “domains” are linked together in geometric arrangements that can be rigid or flexible, depending on the nature of the linker domains. To understand the structure-function relationships in these macromolecules, it would be helpful to visualize the geometric arrangement of domains. Furthermore, accurate spatial representation of domain structure is necessary for coarse-grain models of the multi-molecular interactions that comprise signaling pathways. Here we introduce a new tool, mol2sphere, that transforms the atomistic structure of a macromolecule into a series of linked spheres corresponding to domains. mol2sphere does this with a k-means clustering algorithm. It may be used for visualization or for coarse grain modeling and simulation.

**Availability and implementation:** mol2sphere is available as both a plugin for PyMOL and as a new feature within the SpringSaLaD modeling and simulation software. They run on all desktop platforms and are available with documentation at, respectively, https://nmrbox.org/registry/mol2sphere and http://vcell.org/ssalad. Source code is available for the PyMOL (and python) implementations on the NMRbox platform and for the SpringSaLaD implementation at https://github.com/jmasison/SpringSaLaD.

## Introduction

Biological macromolecules are composed of combinations of distinct sub-molecular domains. The spatial arrangement of these domains and their physical-chemical properties determine the functions of a molecule and how it participates in cell signaling and metabolism. However, accurately representing and visualizing the geometric arrangement of the domain organization in molecules is challenging. All atom representations include too much detail and obscure tertiary and quaternary motifs. The challenge of choosing an appropriate structure representation also presents an issue when simulating dynamics where there is a necessary balance between computational complexity of a model and its fidelity. All-atom parameterizations typical of molecular dynamics simulations are often not computationally practical for spatial modeling of the multi-molecular interactions that govern physiology inside cells. With regard to visualization, cellPACK (1) is an excellent computational tool for representing the shapes of macromolecules and their packing within the crowded environment of the cell. However, it is not designed to visualize the explicit spatial organization of domains or to facilitate coarse grain biophysical models of multi-molecular interactions.

In this work, we introduce a software tool, “mol2sphere”, for representing the domains in a molecule as an interconnected network of spheres. The locations and the sizes of the spheres are derived automatically from the atomic coordinates using a k-means clustering algorithm and may be further refined by the user. The software is implemented as a python module and as a PyMOL (2) plugin, both installed within the NMRbox computational platform (3), which is freely accessible to all members of the academic, government, and not-for-profit communities at https://nmrbox.org. The software is also installed as a utility within the SpringSaLaD simulator (4) which is freely available at http://vcell.org/ssalad. SpringSaLaD uses these coarse grain molecule representations for Langevin dynamics modeling of mutli-molecular reaction-diffusion systems.

### Spherical Decomposition of Macromolecules by k-means Clustering

The objective of the spherical decomposition method is to fill the molecular envelop of a target structure with non-overlapping spheres, thereby providing a coarse representation of the molecule that accounts for both spatial restraints and dynamics. The number of desired spheres is selected as an input parameter and is largely determined by the computational complexity (or visualization granularity) desired. The positions of the sphere centers are determined by k-means clustering of alpha-carbon positions for proteins (by default) or any other atomic subset, depending on molecule type and size. The k-means algorithm partitions the set of atomic positions into k clusters and minimizes the largest separation distance between any atom and the centroid position of its assigned cluster.

The k-means algorithm naturally produces clusters that are “compact” and are thus well-represented as spheres. If the objective is to maximize the volume contained by the spheres and avoid overlap, the radius of each sphere can be set to half the distance to its nearest neighboring sphere center. This greedy approach is simple, whereas a global optimization of sphere radii to maximize the overlap between sphere volumes and molecular volume is a more complex problem that likely yields merely a cosmetic difference, rather than a functional one. One notable exception that benefits from manual intervention is for structures that include an extended linker. The default approach would result in spheres extending well beyond the envelop of the linker, thereby misrepresenting the overall structure in visualizations and removing the flexibility and dynamics from a coarse-grained model that would otherwise be observed. In these cases, it becomes desirable to reduce the sphere radii and/or increase the number of spheres (if computational complexity is not the limiting factor). These manual interventions can be made in the SpringSaLaD interface by specifying closely spaced cluster centers for the k-means algorithm along the linker region, rather than allowing the algorithm to initialize with random centroid positions.

### Running mol2sphere

The implementations of mol2sphere are illustrated with the adaptor protein Nck (5), which contains an SH2 domain and three SH3 domains. It functions to mediate signaling from Receptor Tyrosine Kinases to the actin cytoskeleton (5). Screen shots of the PyMOL plugin and SpringSaLaD utility are shown for the molecule Nck in Figure 1. The atomic coordinates for Nck1 were derived by homology modeling using Modeler (6) and RaptorX (7).

**Figure 1.**
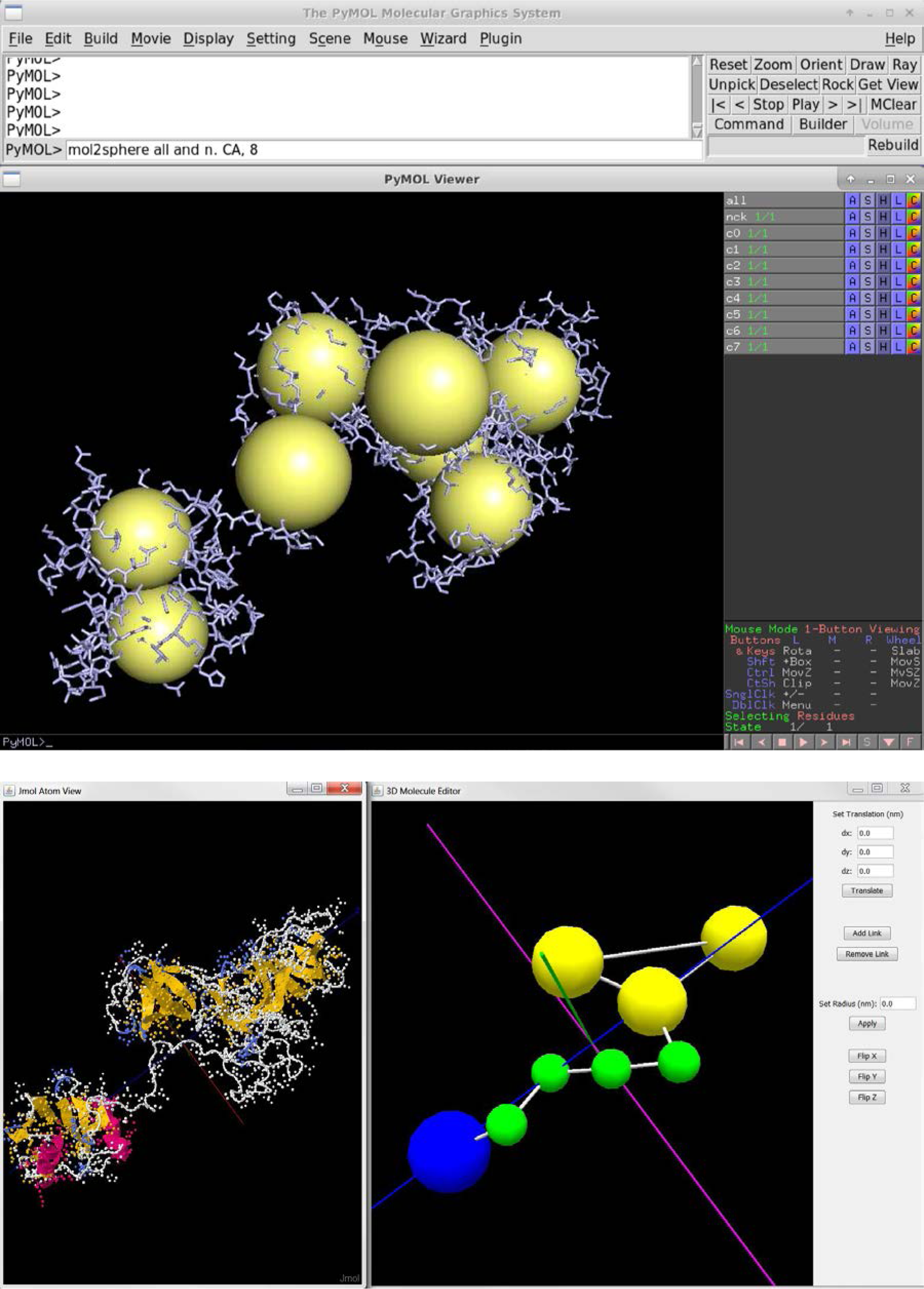
Screenshots of the mol2sphere results in PyMOL (top) and SpringSaLaD (bottom) showing a decomposition of Nck into 8 spheres. The PyMOL example shows the output of the fully automated algorithm. The SpringSaLaD example shows the results of user adjustments to constrain the output to account for known domain and disordered region locations.

The primary way of accessing the mol2sphere utility on the NMRbox platform is through PyMOL, where the utility is pre-loaded, so that it may be invoked in the graphical interface to model and visualize molecules in the PyMOL workspace. At the prompt in PyMOL, the command “mol2sphere” may be issued with no input parameters; this default mode clusters all alpha carbon atoms or all molecules in the workspace into 10 clusters. For greater control, the atom selection (using the native PyMOL selection algebra) and the number of spheres may be passed as input parameters as: “mol2sphere <selection>, <k>”. The utility creates objects in the PyMOL workspace named c0, c1, etc. that are each composed of a single atom at the cluster’s centroid with a van der Waals radius set to the cluster’s radius. Information about the centroid positions, radii, and constituent atoms is printed to the command line and can also be saved to file using PyMOL’s “Save Moleculeߪ” menu. A PyMOL session showing the mol2sphere results for Nck1 are shown in Figure 1, top. The mol2sphere utility is also accessible in NMRbox through a wrapper script on the command line and as a module that may be imported into python scripts. These modes of access do not provide an interactive graphical interface and are provided to facilitate batch processing and integration into other analysis workflows.

In the SpringSaLaD interface, the user is asked to import the .pdb file from the File Menu and then asked how many spheres should be derived from the atomic coordinates (8 in Fig. 1, bottom). The user has the option to allow the k-means clustering algorithm to operate automatically, or to constrain the centers of the spheres by specifying a series of coordinates (in nm units). This option is useful if the positions of the domains of interest are known. This option was utilized in Fig. 1 to create one SH2 domain, four sites in a disordered linker region of the protein and three sites corresponding to the SH3 domains. After the linked sphere structure is created, the user may assign a “site” identity to each sphere with an associated chosen color (blue SH2, green linker sites and yellow SH3 sites). Other options include overriding the radius of a site (the linker sites were reduced in the model of Fig. 1) and adding or removing links (a link was added to constrain the three SH3 spheres into a fixed geometric relationship). As illustrated in Fig. 1, bottom, SpringSaLaD also displays the original atomic structure next to the mol2sphere structure and they can be synchronously rotated for direct comparison; the atomic structure is visualized with Jmol (8) from within SpringSaLaD, so that all the formats for displaying macromolecular structures within Jmol become available. SpringSaLaD can simulate reaction diffusion processes with binding partners for the SH3 and SH2 domains in Nck, where the linked green spheres serve to model a disordered flexible tether. Full instructions can be found on pages 14-17 of the SpringSaLaD v.2 User Guide.

## Acknowledgments

The authors gratefully acknowledge useful discussions with James Schaff at UConn Health. The project is supported by the National Institute of General Medical Science through grants P41 GM103313 and P41GM111135.

